# Pulsed electric fields create pores in the voltage sensors of voltage-gated ion channels

**DOI:** 10.1101/838474

**Authors:** L. Rems, M. A. Kasimova, I. Testa, L. Delemotte

## Abstract

Pulsed electric fields are increasingly used in medicine to transiently increase the cell membrane permeability via electroporation, in order to deliver therapeutic molecules into the cell. One type of events that contributes to this increase in membrane permeability is the formation of pores in the membrane lipid bilayer. However, electrophysiological measurements suggest that membrane proteins are affected as well, particularly voltage-gated ion channels (VGICs). The molecular mechanisms by which the electric field could affects these molecules remain unidentified. In this study we used molecular dynamics (MD) simulations to unravel the molecular events that take place in different VGICs when exposing them to electric fields mimicking electroporation conditions. We show that electric fields induce pores in the voltage-sensor domains (VSDs) of different VGICs, and that these pores form more easily in some channels than in others. We demonstrate that poration is more likely in VSDs that are more hydrated and are electrostatically more favorable for the entry of ions. We further show that pores in VSDs can expand into so-called complex pores, which become stabilized by lipid head-groups. Our results suggest that such complex pores are considerably more stable than conventional lipid pores and their formation can lead to severe unfolding of VSDs from the channel. We anticipate that such VSDs become dysfunctional and unable to respond to changes in transmembrane voltage, which is in agreement with previous electrophyiological measurements showing a decrease in the voltage-dependent transmembrane ionic currents following pulse treatment. Finally, we discuss the possibility of activation of VGICs by submicrosecond-duration pulses. Overall our study reveals a new mechanism of electroporation through membranes containing voltage-gated ion channels.

**Statement of Significance:** Pulsed electric fields are often used for treatment of excitable cells, e.g., for gene delivery into skeletal muscles, ablation of the heart muscle or brain tumors. Voltage-gated ion channels (VGICs) underlie generation and propagation of action potentials in these cells, and consequently are essential for their proper function. Our study reveals the molecular mechanisms by which pulsed electric fields directly affect VGICs and addresses questions that have been previously opened by electrophysiologists. We analyze VGICs’ characteristics, which make them prone for electroporation, including hydration and electrostatic properties. This analysis is easily transferable to other membrane proteins thus opening directions for future investigations. Finally, we propose a mechanism for long-lived membrane permeability following pulse treatment, which to date remains poorly understood.

## Introduction

The integrity of the cell membrane, while essential for life of any biological cell, presents a barrier that needs to be transiently disrupted in order to deliver therapeutic molecules into the cell. High-intensity pulsed electric fields are increasingly used in medicine to achieve such a transient increase in cell membrane permeability. Examples are cancer treatment for enhanced delivery of chemotherapeutic drugs and gene therapy techniques for intracellular delivery of genetic material (1). The applied electric field induces a phenomenon called electroporation or electropermeabilization. Thanks to insights from molecular dynamics (MD) simulations we now understand that one type of events that takes place in the cell membrane is the formation of pores in the membrane lipid bilayer (2, 3).

However, experimental evidence suggests that membrane proteins are affected as well. Particularly voltage-gated ion channels (VGICs) have been identified as targets of the electric field (4). VGICs are a class of transmembrane proteins that respond to changes in the transmembrane voltage (TMV) with conformational rearrangements that lead to opening or closure of an ion-selective pore. They play crucial roles in the generation and propagation of action potentials in electrically excitable cells, including neurons and muscle cells. All VGICs share a common architecture: each of the four protein domains contains six transmembrane segments (S1–S6) and a pore loop between segments S5 and S6. Segments S1–S4 act as the voltage sensor, whereas S5, S6 and the pore loop serve as the pore-forming module (Fig. 1a,b) (5). Segment S4 contains positively charged residues and has the ability to respond to changes in TMV. The movement of S4 then acts on S5 and S6 to open or close the channel pore depending on the direction of the electric field.

**Figure 1:**
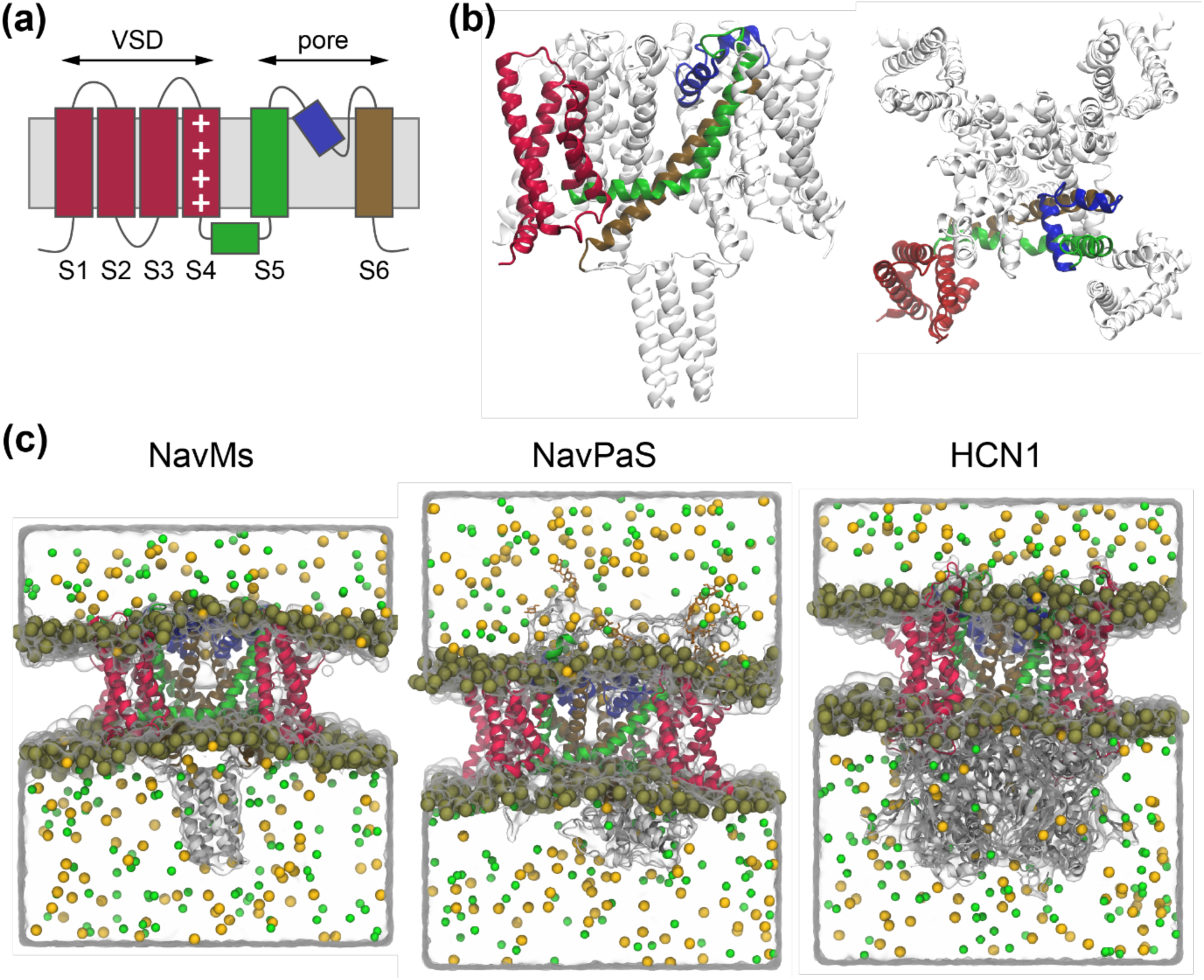
Structure of the voltage-gated ion channels (VGICs) investigated in this study. (a) All VGICs share an overall architecture that includes four subunits, each containing six membrane-spanning segments denoted S1-S6. (b) Side view and extracellular view of the NavMs channel, indicating the segments of a single subunit: VSD (S1-S4) colored in red, S5 together with S4-S5 linker colored in green, pore helix colored in blue and S6 colored in ochre. (c) Simulation box of the investigated systems: NavMs, NavPaS, and HCN1. Ion channels are shown as ribbons colored as in (a). Water molecules are represented as transparent volume, whereas phosphorus atoms of lipid headgroups, Na^+^ ions, and Cl^-^ ions, are shown as gold, yellow, and green spheres, respectively.

Since VGICs are sensitive to changes in TMV and since electroporative pulses induce a TMV of several hundreds of mV, far beyond the physiological resting voltage or voltage generated during action potentials, one can speculate that these channels become perturbed by pulsed electric fields. Indeed, by using advanced patch clamp techniques, electrophysiologists have demonstrated that high-intensity pulses with submicrosecond duration can decrease the ionic currents mediated by different VGICs during action potentials. Nesin et al. (6) studied the effects of 300 or 600 ns pulses on Nav and Cav channels using murine pituitary (GH3) and murine neuroblastoma-rat glioma hybrid (NG108) cells. Electrophysiological measurements revealed a decrease in Nav and Cav currents for pulse amplitudes above 1.5-2 kV/cm. The results from a follow-up study (7) suggested that the decrease in Nav current was not mediated by Na^+^ leakage across the electropermeabilized membrane or downregulation of the Nav channels by a calcium-dependent mechanism. The authors thus proposed as the mechanism either an electroconformational change, or a calcium-independent downregulation of the Nav channels (e.g. caused by alteration of the lipid bilayer). Decrease in Nav channel current was also observed by Yang et al. (8) when they exposed adrenal chromaffin cells to even shorter 5 ns, 50-100 kV/cm pulses. Their analysis suggested that the decrease in Nav current was not due to a change in either the steady-state inactivation or activation of the Nav channels, but instead was associated with a decrease in maximal Na^+^ conductance. This decrease could be observed immediately after the pulse (within 0.5 s, earliest time measured), suggesting a direct effect of the electric field on Nav channels. No effect was observed on Kv channels, whereas a decrease in conductance could also be observed for Cav channels and/or calcium-dependent potassium channels. Although less explored, a decrease in VGICs current has been also reported when using longer millisecond pulses. Chen et al. (9, 10) measured up to ∼40% decrease in Nav and delayed rectifier Kv currents, after subjecting voltage-clamped frog skeletal muscle cells to hyperpolarizing 4 ms, −0.5 V pulses. In all the above-discussed studies the effects depended on the pulse amplitude, with lower amplitude resulting in milder decrease in channel currents. Importantly, all studies also reported that the decrease was not reversible within the observation time (50 min in (10), 10-15 min in (6, 8)).

Apart from decreased current through VGICs, there have been other electric-field effects observed as well. Submicrosecond pulses have been shown to activate specific VGICs (11, 12). However, it remains unclear whether the electric field during such short pulses is able to directly move the VSDs, or whether the channel activation is indirect due to post-pulse membrane depolarization caused by nonselective ion leakage across the electropermeabilized membrane. Furthermore, this activation of VGICs has been found to participate in post-pulse membrane depolarization (13, 14) and intracellular calcium increase (15–17), both of which are important signals that can initiate cell death, proliferation, or differentiation depending on their spatio-temporal profile (18, 19). Understanding the effects of pulsed electric fields on VGICs is thus also interesting for new applications in wound healing and tissue engineering.

Electropermeabilization is associated with a complex set of events including oxidative lipid damage, disruption of the cytoskeleton network and its association with the plasma membrane, and lipid scrambling, which could all affect the function of membrane proteins (2). Yet, the experimental measurements suggested that pulsed electric fields induced some electroconformational change of VGICs during the pulse. Without a molecular-level insight, the mechanisms by which the electric field could affects these structures remain unidentified. In this study we used MD simulations to unravel the molecular events that take place in different VGICs when exposing them to an electric field that mimics electroporation conditions. Since experimental studies consistently reported a decrease in Nav current, we performed simulations of two Nav channels, a bacterial Nav from *Magnetococcus marinus* (NavMs) (20) and an eukaryotic Nav from *Periplaneta americana* (NavPaS) (21). Eukaryotic Nav channels are composed of a single polypeptide containing four homologous but nonidentical domains connected by intracellular linkers; bacterial Navs, on the other hand, are comprised of four identical domains, each being analogous to a single domain of their eukaryotic counterparts, and can thus be used as simple models of eukaryotic channels (22, 23). In addition, we performed simulations of the human hyperpolarization-activated cyclic nucleotide-gated (HCN1) channel (24), which is a nonselective voltage-gated cation channel that is responsible for generation of rhythmic activity in heart and brain. Unlike Nav channels, HCN1 activates under hyperpolarizing TMV. The effects on HCN1 are of interest giving the rapid development of pulsed-electric-field-based cardiac ablation for treatment of heart arrhythmias (25). We found that the three tested channels, NavMs, NavPaS, and HCN1, responded differently to electric pulses, which could be related to structural differences resulting in considerably different hydration and electrostatic profile along their VSDs. Comparing their response thus enabled us to gain an atomistic level insight into the biophysical mechanisms governing their interaction with an electric field and to relate their propensity to be porated with their biophysical characteristics.

## Methods

### Systems preparation

The computational systems with NavMs (PDB 5HVX), NavPaS (PDB 5X0M), and HCN1 (PDB 5U6O) were built using CHARMM-GUI web-server (26). Briefly, each protein was embedded into a 1-palmytoyl-2-oleoyl-phosphatidylcholine bilayer and solvated with 150 mM NaCl solution. CHARMM36 force field (27, 28) was used for proteins, lipids, and ions, and TIP3P model (29) for water. The composition of each system is reported in Table S1 in the Supporting Material.

### MD simulations

All simulations were performed in GROMACS 2016 (30, 31). Each system was first minimized using the steepest descent algorithm. The equilibration/production run was then carried out using leap-frog integrator with a time step of 2.0 fs, Nose-Hoover thermostat (τ = 0.4 ps, T = 300 K) (32, 33) and Parrinello-Rahman barostat (τ = 5 ps, P = 1 bar, semi-isotropic coupling) (34, 35). During the first 100 ns the protein atoms were restrained to their initial positions, after which the simulations were continued for 400-1200 ns without restraints (400 ns for NavMs; 1200 ns for NavPas; and 700 ns for HCN1). The long-range electrostatic interactions were calculated using Particle Mesh Ewald method (36) together with a Fourier grid spacing of 0.15 nm and a cutoff of 1.2 nm. A switching function was used between 0.8 and 1.2 nm to smoothly bring the short-range electrostatic interactions and the van der Waals interactions to 0 at 1.2 nm. The chemical bonds were constrained to their equilibrium values using the LINCS algorithm (37). Periodic boundary conditions were applied.

To simulate the exposure to an electric pulse, we added a force *qE*_*z*_ to every atom carrying a charge *q* (2, 3). This method induces a TMV that is approximately equal to the product of the imposed electric field and the simulation box length, TMV = *E*_*z*_ *L*_*z*_. The *E*_*z*_ was chosen based on the following considerations. Previous simulations on pure lipid bilayers showed that formation of a lipid pore occurs faster with increasing electric field magnitude; in order to observe lipid pores within few tens of nanoseconds, simulations have been typically conducted by applying an electric field resulting in TMV above ∼3 V (42, 43). Experimentally, however, the TMV that can be built on the membrane is limited, as the membrane starts discharging through pores after it becomes electroporated. According to measurements, performed using pulses with duration of 60 ns and longer together with voltage-sensitive dyes (44, 45) and microelectrodes (46), the TMV of ∼1.5 V is the largest that a cell membrane can sustain before discharging. Thus, we chose the value of the electric field such that it resulted in the TMV of about ±1.5 V. The chosen value is a compromise between trying to observe an effect of the electric field within reasonable simulation time and at the same time staying within realistic experimental TMV values. These simulations were performed in NVT ensemble (constant number of atoms, volume, and temperature), which kept the simulation box size constant. This ensured that the voltage applied across the membrane remained constant.

We also performed two additional control simulations on NavMs channel by using an electric field in NPT ensemble, and applying a constant charge imbalance (38) (Section S2). In the charge imbalance method, the TMV is built by (i) separating the water bath into two parts by adding an additional lipid bilayer or a vacuum layer, and (ii) transferring a certain number of positive and negative ions across the membrane, which creates the required charge imbalance. The charge imbalance is kept constant despite the transport of ions through transmembrane pores throughout the simulation by replenishing the charge imbalance using a Monte Carlo setup (39).

Simulations in which we characterized the properties of pores formed in VSDs, including their ionic conduction at 3x lower electric field and their stability in the absence of an applied electric field, were carried out in the NPT ensemble.

### Analysis

Trajectories were visualized with VMD (40). Ion passage through pores formed in the system was determined by a custom Matlab code based on the positions of ions extracted from the trajectories (see also Section S3). The code is available at https://github.com/delemottelab. Free energy profiles of water molecules along VSDs were determined based on kernel density estimate of the probability distribution of the positions of water molecules extracted from the trajectories at 0 V (see also Section S4). To calculate electrostatic profiles along the VSDs, additional 2-ns-long simulations under an electric field were performed while keeping the position of the protein heavy atoms restrained. These short simulations were carried out starting from 10 different configurations extracted from the last 100 ns of the equilibration trajectories. For each of these short trajectories we determined the 3D electrostatic potential using VMD tool PMEpot (options: ewaldfactor 0.5, grid 1.0 Å, padding along z direction of ∼15 nm) (41). From each of the ten 3D profiles we determined a 1D electrostatic profile along the VSD by averaging the potential along a cylinder with a diameter of 0.5 nm centered at the center of mass of the VSD. Finally, we computed the average and standard deviation of the ten 1D profiles.

## Results

When a cell is exposed to an electric field, the part of its membrane facing the positive electrode becomes hyperpolarized while the part facing the negative electrode becomes depolarized (see the scheme in Fig. 2a). Thus, for each of the three investigated channels (NavMs, NavPaS, and HCN1) we generated up to 600-ns-long trajectories under hyperpolarizing and depolarizing TMV. The simulations were run until a conducting aqueous pathway was formed somewhere in the system, which took about 100-500 ns. Such pathway could be formed either in the middle of a VSD or in the lipid bilayer surrounding the channel. The types of pathways that we observed are schematically depicted in Fig. 2b. The graphs in Fig. 2a summarize the results obtained for all channels, where different markers show the time of the first ion passage, either through a VSD-associated conductive pathway or through a lipid pore (see figure legend). The horizontal bars in Fig. 2a indicate the time at which we stopped the simulation. More detailed description of individual simulations is given in the following sections.

**Figure 2:**
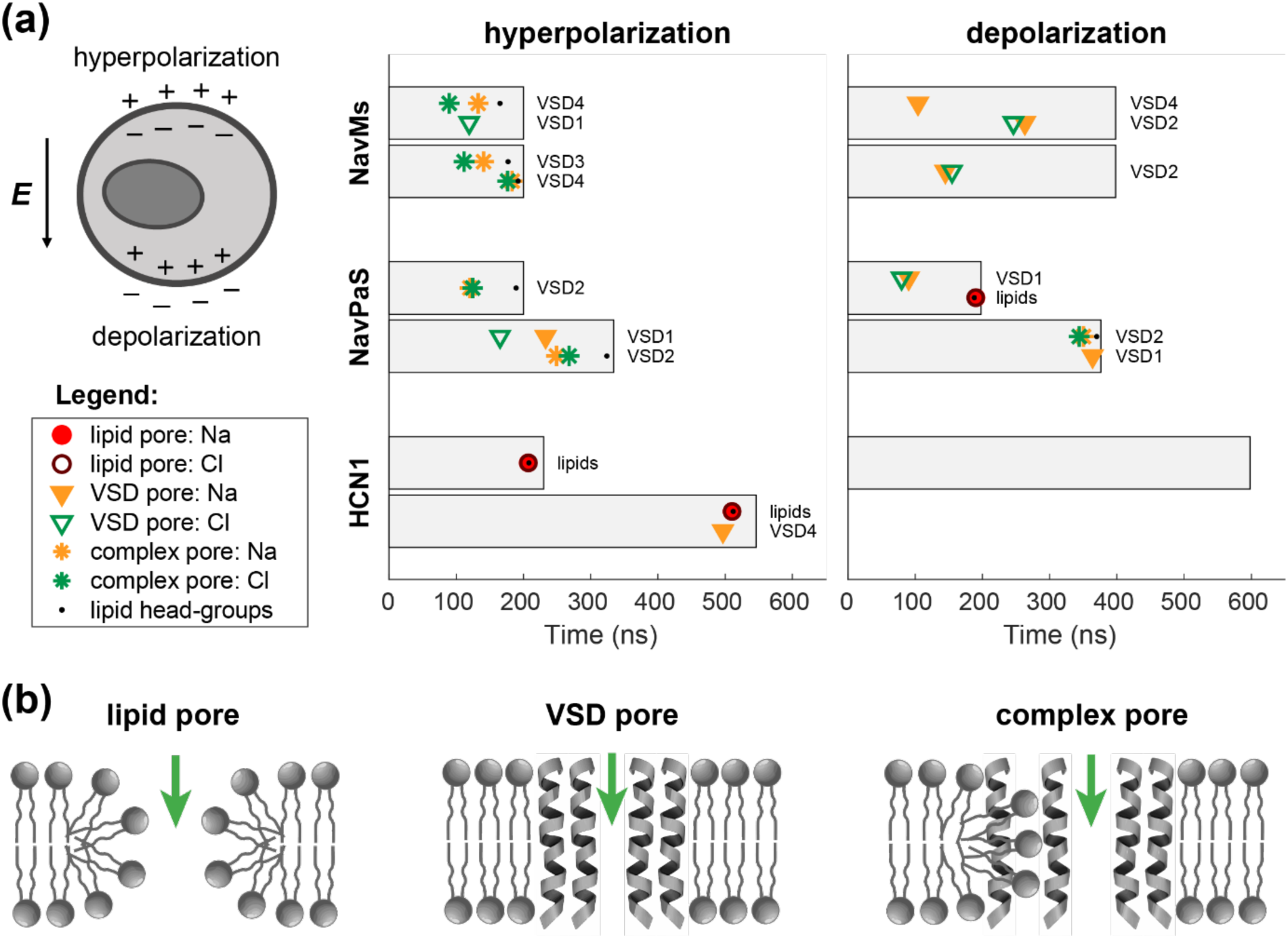
Summary of the simulation results. (a) Left: Scheme showing how ions redistribute around the cell membrane upon exposure to an electric field; on the side facing the positive electrode (anode) the membrane becomes hyperpolarized, whereas on the opposite side it becomes depolarized. Right: Graphs show the time of the first Na^+^ and Cl^-^ ion passages, either through a lipid pore or through a VSD-associated conductive pathway, as indicated by different markers. Black dots denote the time at which a pore became stabilized by lipid head-groups (i.e., the time when at least two lipid head-groups were found within 0.5 nm from the middle of the membrane). The horizontal bars correspond to the simulation length. The text next to each horizontal bar gives the index of the VSD in which the conductive pathway was formed. The data presented in this figure is also tabulated in Table S3. (b) Schematic representation of the types of conductive pathways observed in simulations.

### Electroporation of the NavMs channel

We first performed a simulation of NavMs channel under a hyperpolarizing TMV of 1.5 V. The VSDs of NavMs are already hydrated in the absence of an external electric field (Fig. 3a). After the onset of the electric field, more water molecules entered the VSDs (Fig. 3b). In one of these domains denoted as VSD4 the S3 helix moved away from S1, S2 and S4, which further promoted hydration of the domain’s interior (Fig. 3c). Compared to the other helices, S3 is weakly connected to the rest of the voltage sensor: first, it is a short helix, which limits the number of interactions it can form, and second, it has only one salt bridge interaction with S4 unlike, for instance, S2 that has two of such salt bridges; we anticipate that this weak connection resulted in the S3 helix to be detached first under an excessive electric field. Shortly after S3 moved, at ∼90 ns VSD4 was crossed by the first Cl^-^ ion (Fig. 3d), which was followed by the passage of the first Na^+^ ion at ∼130 ns. We will further refer to this conductive pathway as the “VSD pore”. Along the simulation this VSD pore expanded and eventually became stabilized by lipid headgroups that migrated towards the middle of the membrane (Fig. 3e-h). Following previous suggestions in the literature (47, 48) we will refer to this lipid-stabilized structure as the “complex pore”. With expansion of the complex pore, the S4 helix lost its secondary structure, and VSD4 began to unfold, as shown in Figs. 3g-i and Movie S1 and Movie S2. In this simulation, another VSD pore was formed in VSD1 at ∼120 ns, which allowed the passage of a single Cl^-^ ion.

**Figure 3:**
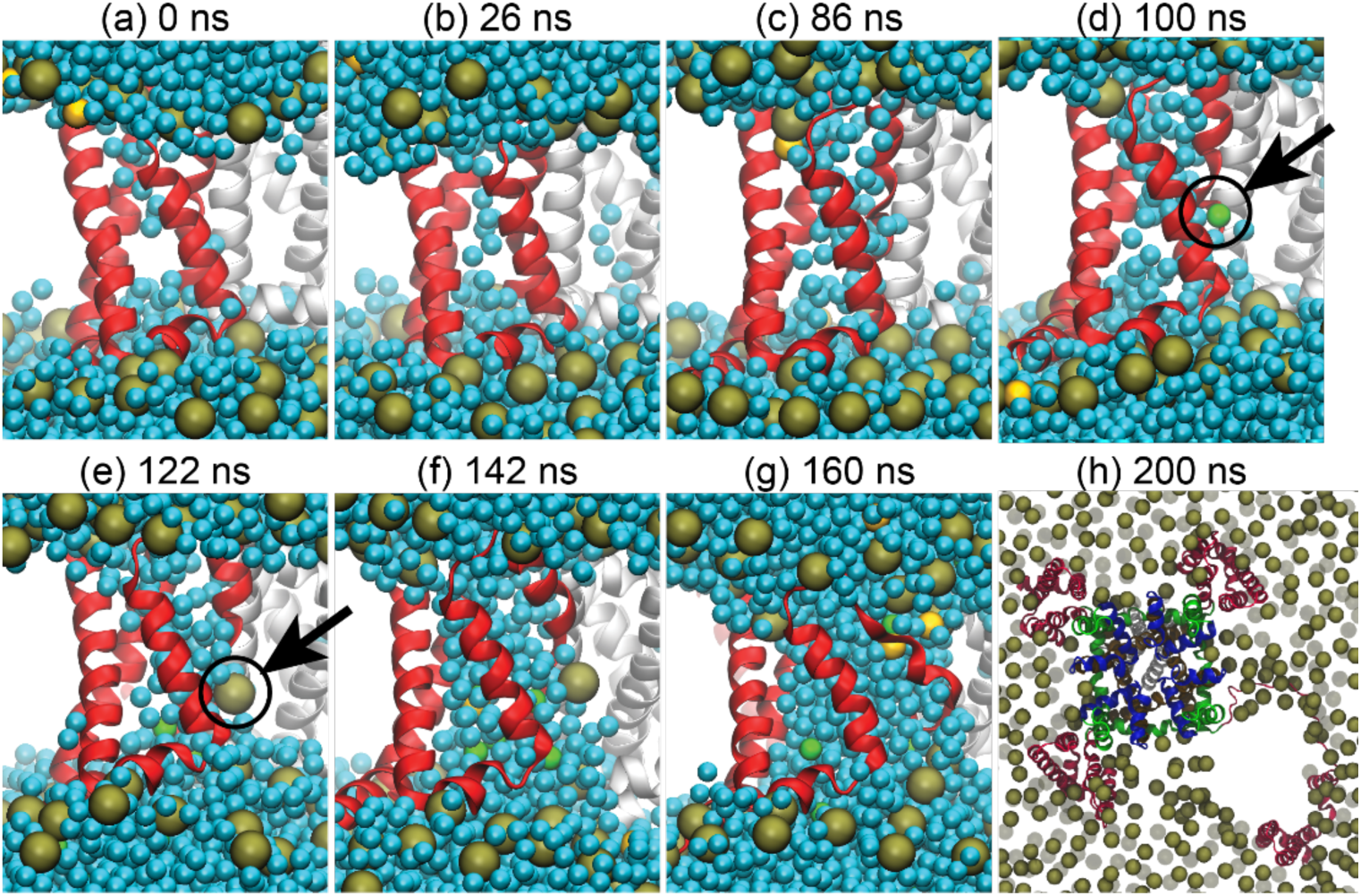
Formation of a complex pore in VSD4 of NavMs channel. The first Cl ion passed through the VSD at ∼90 ns after the onset of the electric field. More water and ions entered the VSD and this VSD pore became stabilized by lipid headgroups forming the so-called complex pore. As the complex pore expanded, the VSD began to unfold from the channel. The last image at 200 ns shows the unfolded VSD when viewing the channel from the extracellular side. The VSD is colored in red, water in cyan, lipid phosphorus atoms in gold, and sodium and chloride ions in yellow and green, respectively. Black arrows mark the first Cl ion within VSD and the first lipid headgroup moving into the pore.

To verify that these results are reproducible, we performed a second 200-ns-long simulation under the same conditions. Again, we observed formation of a complex pore, but this time in VSD3, with the first passage of a Cl^-^ ion occurring at ∼110 ns. Subsequently, a second complex pore was formed in VSD4, suggesting that such pores can be formed in the same voltage-gated ion channel. The NavMs has four identical VSDs, therefore it is expected that pores can be formed randomly in any of them with equal probability.

We further performed simulations under depolarizing TMV of −1.5 V. In this case we also observed formation of VSD pores, but they didn’t expand into complex ones on the 400 ns timescale. In the first out of two simulations, VSD pores were formed in VSD2 and VSD4. In the second simulation, a single Na^+^ and a single Cl^-^ ion passed through VSD2, confirming that formation of complex pores is more difficult under depolarizing compared to hyperpolarizing TMV.

In all four simulations we also observed passage of Na^+^ through the central pore of the channel (Fig. 4). The structure of NavMs is presumably in an open state (20), although in simulations we observed that the channel pore became fully hydrated and conductive only upon application of a strong (beyond physiological values) electric field. Furthermore, the pore stopped conducting as soon as we either lowered the electric field three-fold or completely turned it off, although in both cases the channel pore still became fully hydrated occasionally, similarly as already shown in Fig. 4a at 2 ns. By counting the number of Na^+^ ions that passed through the channel pore within ∼150 ns at ±1.5 V, we estimated its conductance to be ∼17 pS and ∼2 pS in the two simulations under hyperpolarizing TMV, and ∼17 pS and ∼15 pS in the simulations under depolarizing TMV (all at 27°C). The estimated conductance appears lower than the experimental value (33 pS at 22°C) (49). We therefore speculate that the channel structure that we used in our simulations does not correspond to a fully open state and that the passage of Na^+^ ions was only enabled by the high electric field, which promoted the hydration of the hydrophobic gate along the channel pore.

**Figure 4:**
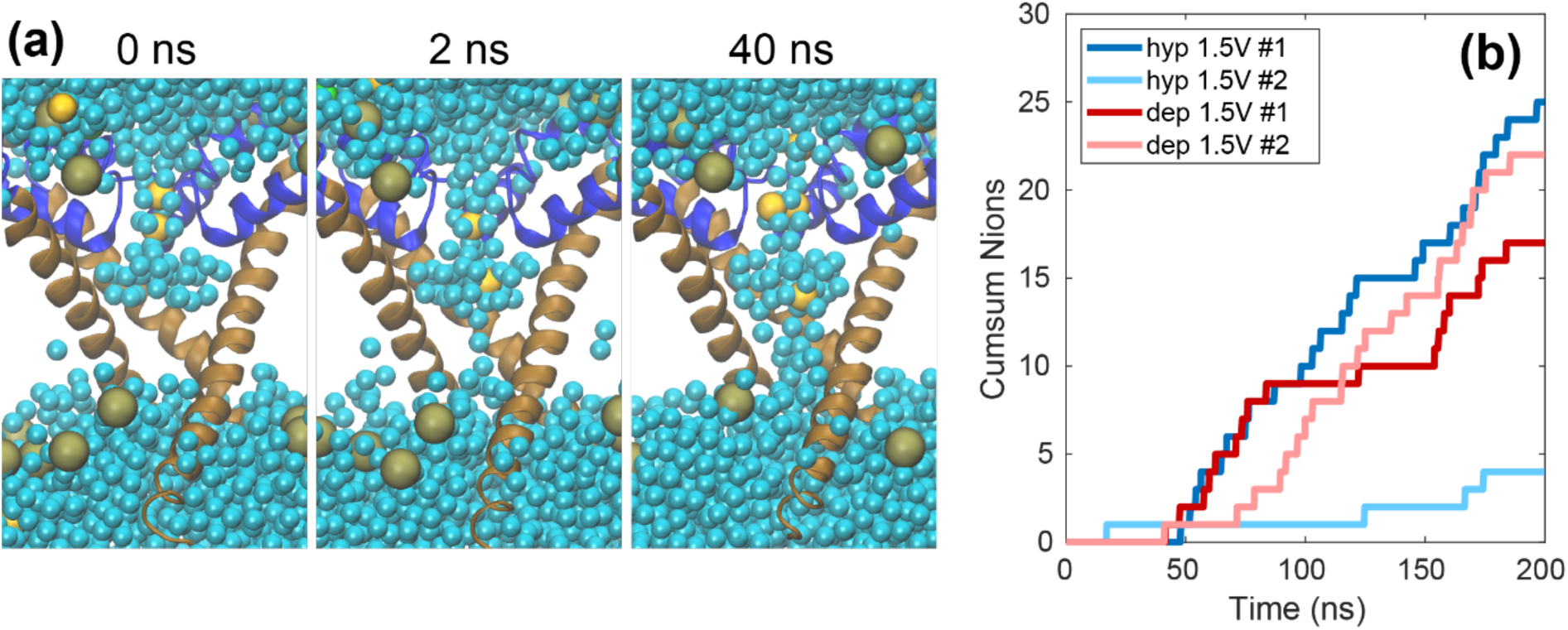
Passage of Na+ ions through the central pore of the NavMs channel. (a) Before electric field application, the bottom half of the pore is dehydrated. After the application, a stable water bridge is formed in the bottom half of the channel pore and the number of water molecules hydrating the bottom half gradually increases, allowing the transport of Na+ ions. (b) Cumulative sum of the number of ions that passed through the channel pore in simulations under hyperpolarizing (blue shades) and depolarizing TMV (red shades).

### Electroporation of the NavPaS channel

To verify that the formation of VSD pores and complex pores is not observable only in NavMs, we studied also the eukaryotic NavPaS channel. As for NavMs, we performed two simulations for NavPaS under hyperpolarizing and two under depolarizing TMV. In the first simulation under hyperpolarizing TMV a VSD pore was formed in VSD2 at ∼120 ns, which then expanded into a complex pore. In the second simulation, VSD pores were formed in VSD1 and VSD2 at ∼233 ns and ∼249 ns, respectively. While the pore in VSD1 did not expand considerably and mainly enabled the transport of Cl^-^ ions, that in VSD2 expanded into a complex pore (Movie S3 and Movie S4). In the first simulation under depolarizing field, a VSD pore was formed in VSD1 at ∼80 ns. In addition, at ∼190 ns a lipid pore was formed in the lipid bilayer surrounding the channel, as illustrated in Fig. 5. In the second simulation, a VSD pore was again formed in VSD1 and also in VSD2, but only in VSD2 the pore expanded into a complex one. Importantly, unlike NavMs, NavPaS has four different VSDs, therefore differences in their perturbation are expected.

**Figure 5:**
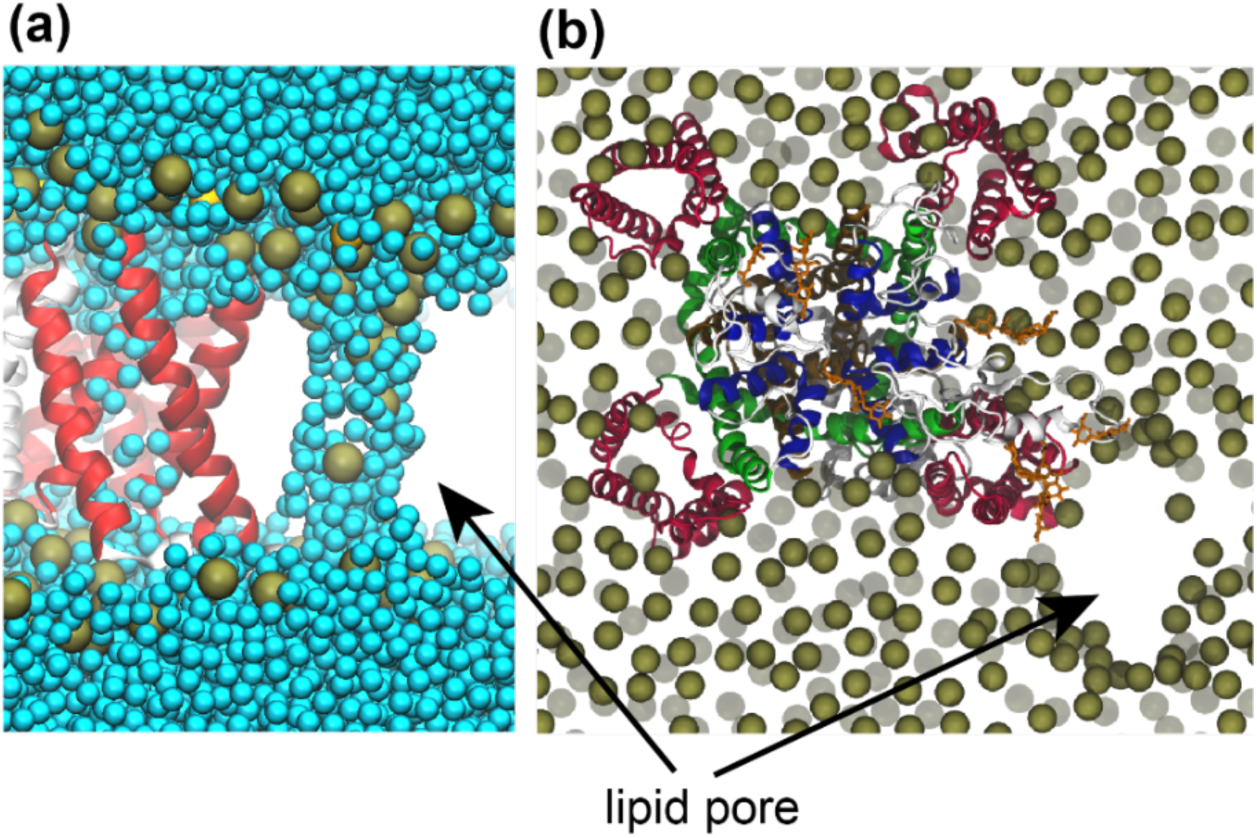
Formation of a lipid pore in the NavPaS system. (a) Snapshot showing formation of a lipid pore close to, but not associated with, the protein. Representation of atoms is similar to Fig. 3. (b) Top view of the lipid pore. Representation of atoms is similar to Fig. 1.

Ion passage through the central pore was not observed in any of the simulations. It should be noted that the functional state of the NavPas structure is unclear, since it was not possible to electrophysiologically characterize this channel (21).

### Electroporation of the HCN1 channel

In addition to Nav channels we performed simulations on HCN1. The VSDs of HCN1 turned out to be much less prone to poration than those of NavMs and NavPaS (Fig. 2). In one out of two simulations performed under hyperpolarizing TMV we observed the passage of two Na^+^ ions through VSD4 at ∼497 ns and ∼525 ns. However, in both simulations we also observed formation of a lipid pore in the lipid bilayer surrounding the protein, suggesting that formation of pores within the identical VSDs is energetically less favorable than that of lipid pores (Movie S5 and Movie S6). In the single simulation performed at depolarizing TMV we observed no pores within the entire 600 ns run. The structure of this channels is expected to be in the closed state, and in none of the simulations we observed passage of ions through the channel pore.

### VSD pores and complex pores form more easily in VSDs which are more hydrated and are electrostatically more favorable for the entry of ions

The results presented above showed that it is possible to observe conduction of ions through one or more VSDs in all three channels. However not all VSDs are equally likely to be perturbed by the electric field (Fig. 2). Considering all simulations performed, we observed ionic conduction through all of the identical VSDs of NavMs. In addition, in NavMs we observed the expansion of VSD pores into complex ones in all VSDs except VSD1 (see also the control simulations reported in Table S2). In NavPaS, which has four different VSDs, we observed ionic conduction only through VSD1 and VSD2. Moreover, VSD2 formed a complex pore, whereas VSD1 did not. In HCN1, formation of a lipid pore was more favorable compared to a VSD pore, as we could only observe the passage of two sodium ions through VSD4 in one out of the three simulations, while a lipid pore was formed in two of these simulations. Overall, these differences suggest that there should be some features of the VSD, which make it more prone to porate. We thus hypothesized that VSD pores and complex pores are formed more easily in VSDs which are more hydrated. This hypothesis is based on previous MD simulations of pure lipid bilayers showing the crucial role of water molecules in formation of lipid pores (50), as well as on the observations that in a lipid bilayer a pore is formed more easily, if this bilayer is pre-embedded with water molecules (51).

To investigate the hydration profile of individual VSDs, we estimated the free energy of water molecules as a function of their position along the VSDs, by analyzing the probability distribution of these molecules in the absence of an external electric field (Fig. 6a). The free energy profiles indeed show the lowest barriers in the VSDs where we observed formation of VSD pores and complex pores, i.e. in the VSDs of NavMs and in VSD1 and VSD2 of NavPaS. The water probability distribution in the two other VSDs of NavPaS and the four VSDs of HCN1 have hydrophobic gaps resulting in considerably higher free energy barriers (see also 2D images showing the position of water molecules in Fig. 6c and Section 4). Furthermore, for all VSDs from all the channels we plotted the height of the free energy barrier versus the time of the first ion passage (either Na^+^ or Cl^-^) through the VSD. The graph in Fig. 6b shows that there is a positive correlation between these two variables, confirming that hydration of the VSD is an important feature, which contributes to VSD’s propensity for poration. To corroborate this hypothesis further, we made three mutations along the S1 helix of the HCN1 VSDs, where we replaced three nonpolar residues with a polar serine (52). The mutations increased the hydration of the VSDs, and decreased the free energy barrier for water molecules, as shown in Fig. 6a. We then performed simulations of the mutant HCN1 under hyperpolarizing TMV. In one out of two simulations a VSD pore was formed at ∼250 ns after the onset of the electric field, followed by its transformation into a complex pore; this observation indicates that the VSDs of the mutant HCN1 can be more easily porated compared to the wild type channel and goes in line with our hypothesis (Table S3 and Fig. S8).

**Figure 6:**
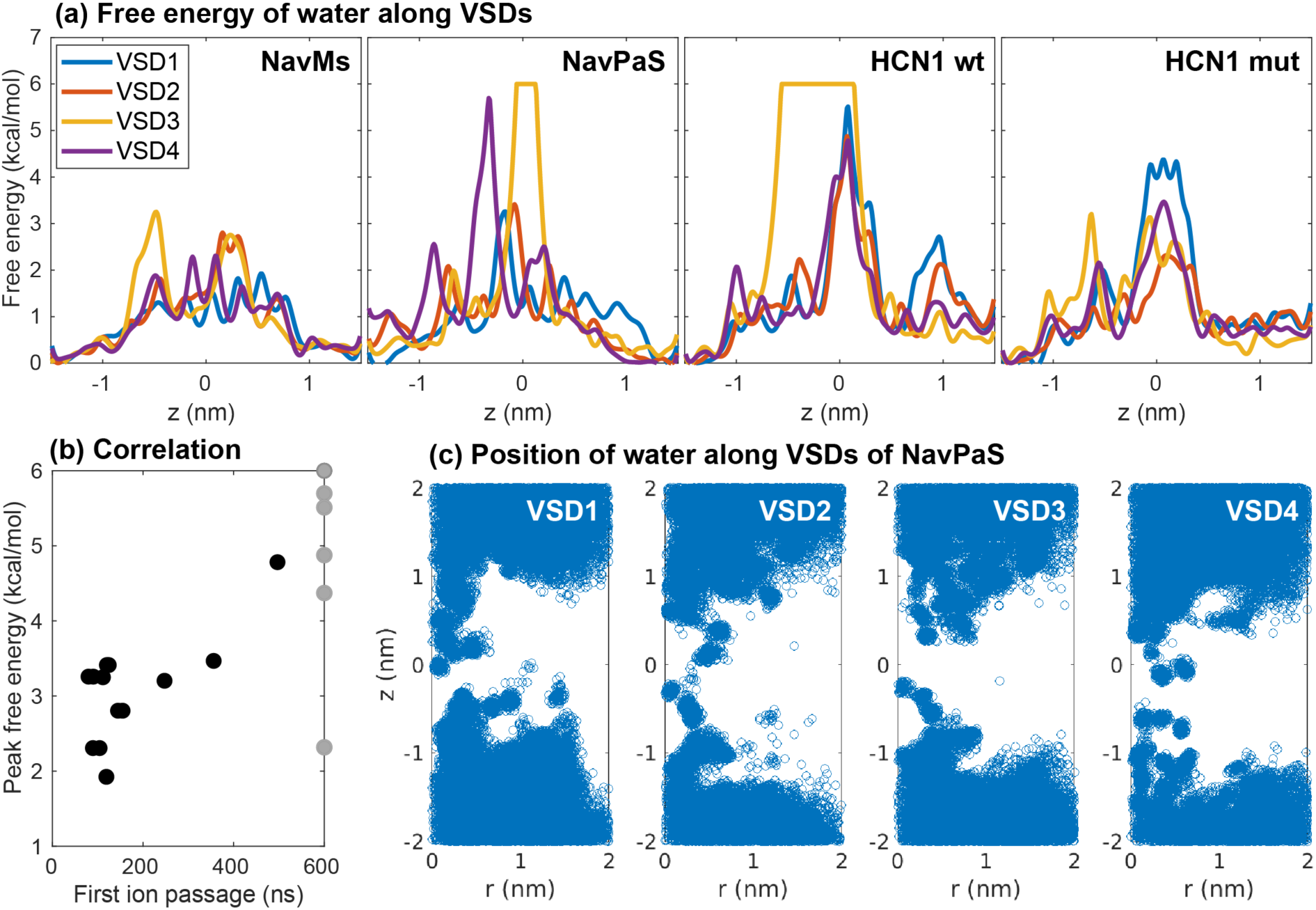
Hydration of VSDs. (a) Free energy profiles of water permeation through VSDs principal axis (z) of NavMs, NavPaS, wild-type HCN1, and mutant HCN1, averaged over 200 ns. Note that the free energy barriers are cut at 6 kcal/mol. (b) Correlation between the height of the free energy barrier and the time of the first ion passage (either Na+ or Cl-) through a VSD. Ion passage times are taken from all simulations reported in Table S3. Gray dots represent VSDs that were not porated in any of the simulations. The outlier at (600 ns, 2.3 kcal/mol) corresponds to VSD2 of HCN1 mutant; note that we performed only two simulations for this channel. (c) Positions of water molecules in each VSD of NavPaS projected along the VSD radius (r) and the VSD principal axis (z). The VSDs’ center of mass is located at (0,0). Blue circles show all positions extracted from 200 frames of the last 200 ns of the equilibration trajectory.

The electrostatic potential inside the VSD is another candidate feature that contributes to VSD’s propensity for poration. The simulations showed that in NavMs complex pores could only be formed under hyperpolarizing TMV. In NavPaS we observed formation of a complex pore in VSD2 regardless of the TMV polarity, whereas in VSD1 we only observed formation of a VSD pore. Moreover, the pore in VSD1 was considerably more selective for Cl^-^ ions under depolarizing TMV, allowing the passage of 9 Cl^-^ compared to only 1 Na^+^ ion (see Table S3). We hypothesized that the observed asymmetry in pore formation and conduction is due to the asymmetric distribution of charges along the VSDs. Fig. 7 compares the electrostatic potential profiles along one of the VSDs of NavMs (all four VSDs show a similar profile, Fig. S14) and VSD1 and VSD2 of NavPaS. Note that the electrostatic potential was determined under an applied electric field before poration. The NavMs VSD has a high positive peak at the extracellular side, which attracts Cl^-^ ions but repels Na^+^. The transport of ions is thus impeded under depolarizing TMV, for which we observed formation of VSD pores, but not complex pores. In VSD2 of NavPaS, in which a complex pore was formed under both hyperpolarizing and depolarizing TMV, the electrostatic profile is similar at either side of the membrane. In VSD1 of NavPaS the electrostatic potential also shows a positive peak at the extracellular side. Accordingly, we observed selective conduction of Cl^-^ ions through this VSD under depolarizing TMV, allowing passage of 9 Cl^-^ compared to only 1 Na^+^ ion (see Table S3). Interestingly, the height of the peaks for VSD1 and VSD2 are similar, yet complex pores were formed only in VSD2. This suggests that there are additional features of the VSDs that influence pore formation such as, for instance, the salt-bridge connections. VSD2 has fewer salt-bridge connections between S1-S4 helices than VSD1 (3 vs. 4, see Table S4), which could be the reason why it unfolds more easily.

**Figure 7:**
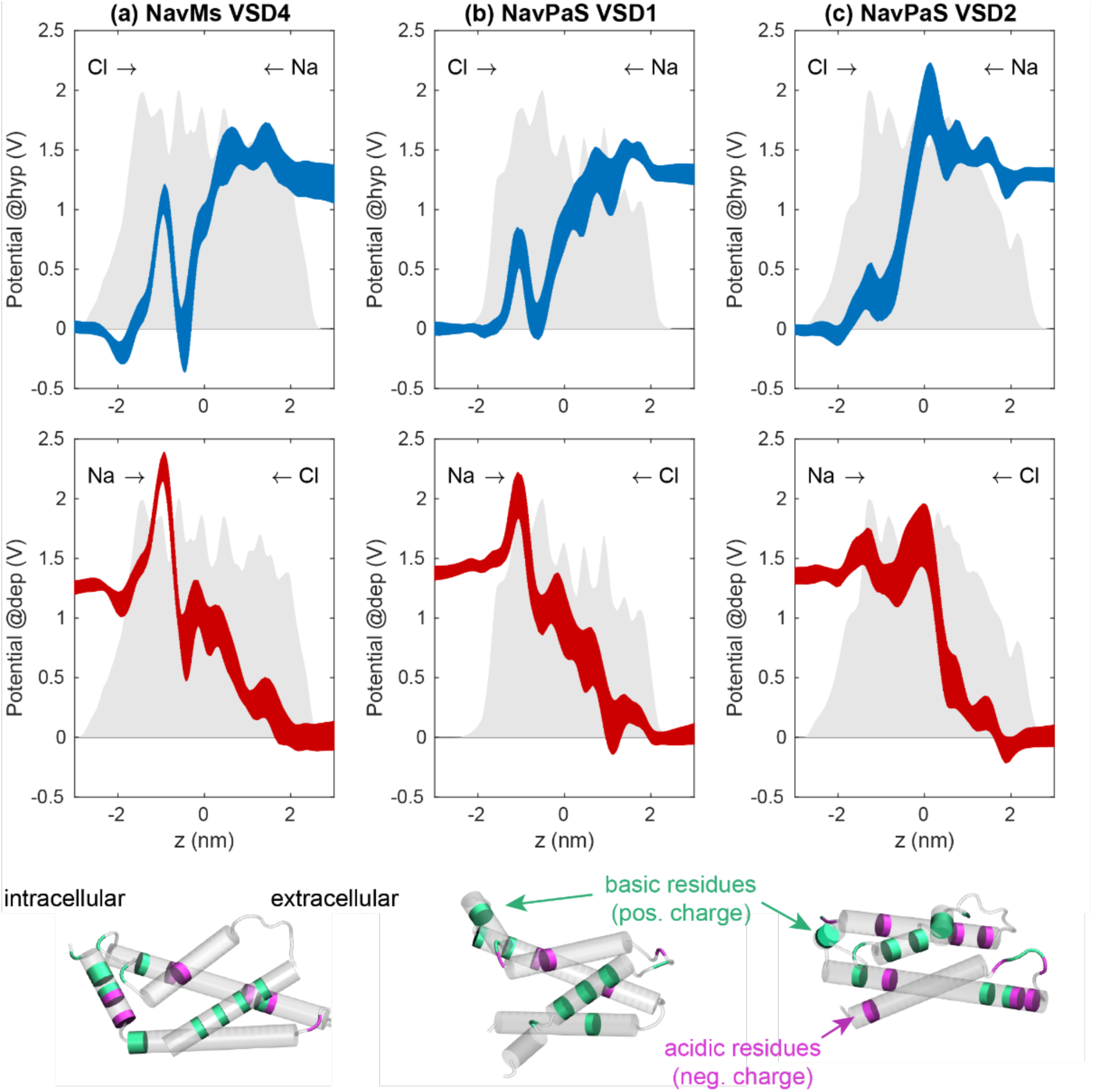
Electrostatic profiles along different VSDs under hyperpolarizing (blue lines) and depolarizing TMV (red lines). (a) VSD4 of NavMs. (b) VSD1 of NavPaS. (c) VSD2 of NavPaS. The line thickness corresponds to standard deviation of ten 1D profiles. Grey area at the back of each graph shows the mass density profile of the VSD. Images below graphs show the position of charged residues on each VSD: positively charged ARG, HIS, and LYS are colored green, negatively charged residues ASP and GLU are colored magenta. Intracellular side is on the left.

### Complex pores can conduct ions preferentially in one direction

To investigate further the asymmetric ion conduction through complex pores, we performed additional simulations of NavMs and NavPaS with these pores under 3x lower electric field, resulting in TMV ≈ *E*_*z*_ *L*_*z*_ = 0.5 V. We reduced the electric field to prevent the complex pores from further expansion, and we characterized ionic transport through such stabilized pores (53, 54). As expected, the conduction of ions through the complex pore was considerably larger under hyperpolarizing TMV, especially for Cl^-^ ions, whereas the conduction through the complex pore in VSD2 of NavPaS was less dependent on TMV polarity (Fig. S16). More specifically, within 100 ns of the simulation, the NavMs complex pore was crossed by 37 Na^+^ and 195 Cl^-^ ions under hyperpolarization, and by 20 Na^+^ and 52 Cl^-^ ions under depolarization. The NavPaS complex pore was crossed by 104 Na^+^ and 276 Cl^-^ ions under hyperpolarization, and by 99 Na^+^ and 189 Cl^-^ ions under depolarization. Both complex pores were less conductive to Na^+^ compared to Cl^-^ ions. Similar result has been observed for lipid pores, where the selectivity could be explained by lower bulk mobility of Na^+^ ions and their binding to lipid head-groups, which affects the electrostatic environment inside the pore (55).

### Complex pores are more stable than lipid pores

To investigate the post-pulse stability of complex pores, we conducted a 1-microsecond-long simulation of the NavMs complex pore, presented in Fig. 3, in the absence of an electric field. Fig. 8a shows the configuration of the pore at 0 ns, 100 ns, and 1000 ns after switching off the electric field. The pore reduced in size but remained stabilized by lipid headgroups even after 1000 ns. Ions were able to enter and pass through the pore by diffusion (Fig. 8b).

**Figure 8:**
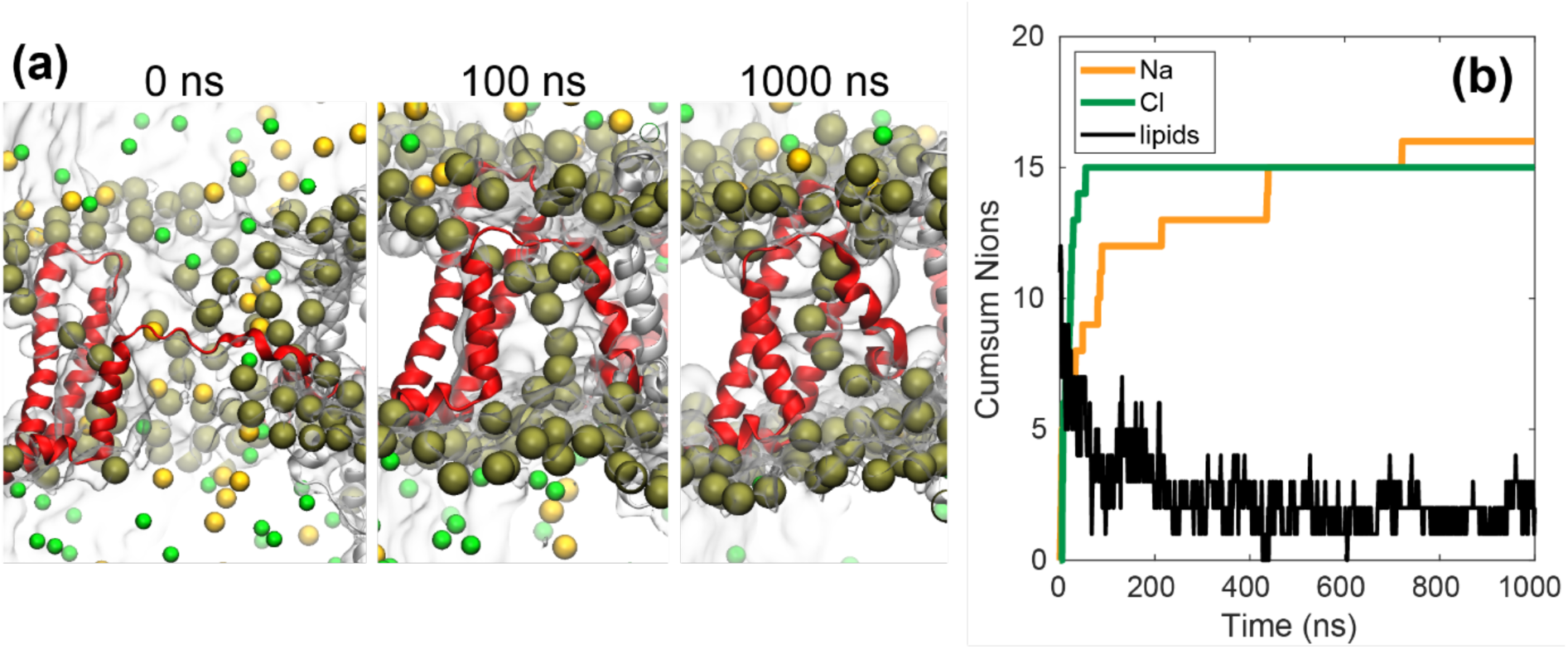
Post-pulse stability of a complex pore. (a) Configuration of a complex pore in NavMs at 0, 100, and 1000 ns after turning off the electric field. Representation of atoms is the same as in Fig. 3; water is represented as a transparent surface. (b) Cumulative number of ions that passed through the complex pore after turning of the electric field. The yellow and green lines show the cumulative sum of the number of Na^+^ and Cl^-^ that passed through the VSD pore, respectively; the passage of ions is here mediated by diffusion and can occur either from intracellular to extracellular side or vice versa. The black line shows the number of lipid phosphorus atoms that are close to the VSD pore and within 0.5 nm of the z-position of the lipid bilayer’s center of mass. The number of lipids that stabilize the VSD pore decreases with time but stabilizes above zero, meaning that even at 1 microsecond after turning off the electric field the VSD pore is still stabilized by lipids.

### Salt bridges reorganization in VSDs

The ability of VSDs to respond to changes in TMV, is granted by positively charged residues of the S4 segment. Being embedded into a low dielectric medium these residues are primary elements to sense an applied electric field. Inside a VSD they interact with negative counterparts coming from the remaining S1-S3 segments through salt bridges. Upon electric field application, the S4 residues move along its direction causing the disruption of existing salt bridges and formation of new ones (56–58). In our simulations, application of the electric field modified the salt-bridge connections within the investigated VSD: some existing salt bridges were broken, and new salt bridges were formed. An example is depicted in Fig. 9a and Movie S7, which shows how salt bridges were perturbed in VSD2 of NavMs channel under hyperpolarizing TMV (note that VSD2 was not porated in this simulation). Before application of the electric field, there were four connections formed by positively charged arginine residues on the S4 helix and negatively charged residues on the S1-S3 helices (Fig. 9a). Upon an electric field application, all these salt bridges were broken and ARG106 shifted such that it formed a new connection with ASP81. After turning off the electric field this new connection ARG106-ASP81 remained stable during at least 1 μs (last time tested). Two other connections, ARG103-ASP49 and ARG109-ASP81, that existed before the electric field application, were reformed (Figs. S17-S20). Similar breakage and formation of salt bridges was also observed in other NavMs VSDs and in other channels, as shown in Tables S4-S6. It is interesting to note that the entry of ions into the VSDs was not necessary for the breakage of salt bridges, but it could facilitate this breakage, as shown in Fig. 9b.

**Figure 9:**
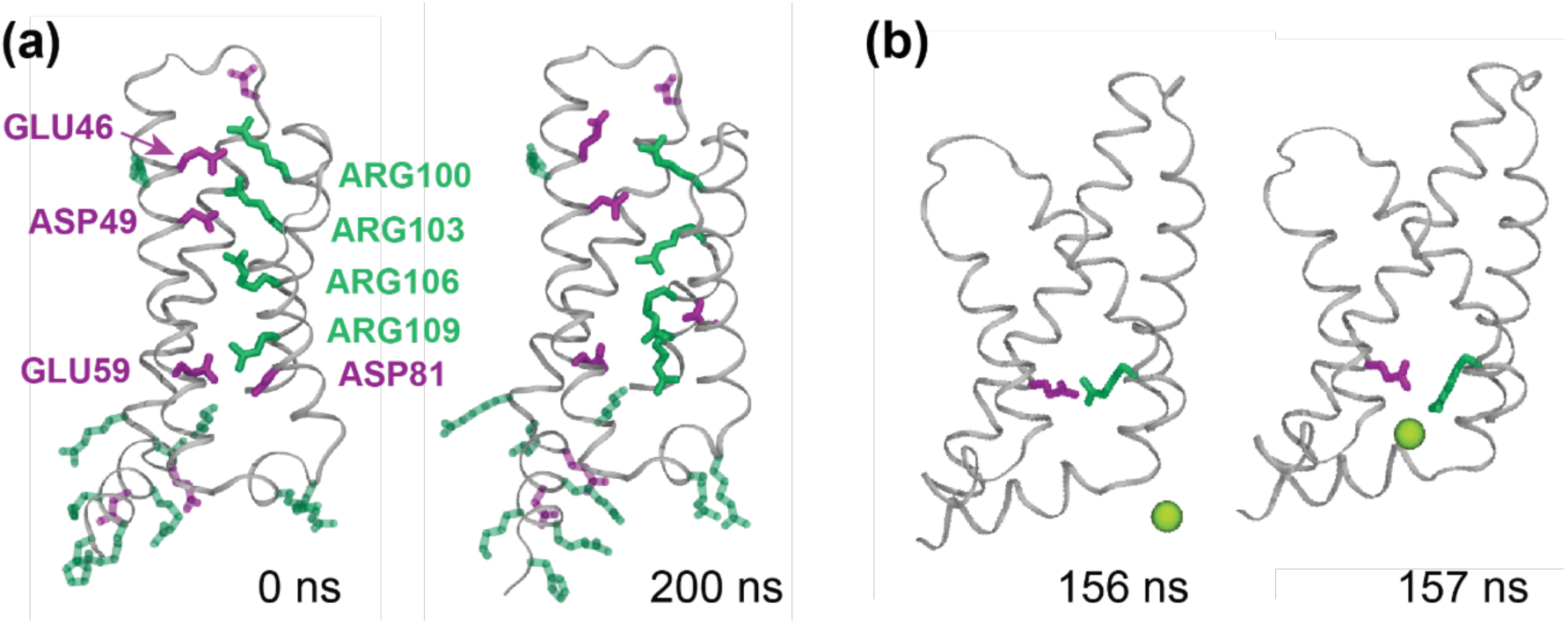
Reorganization of salt bridges in VSDs. (a) Example from the first simulation of NavMs under hyperpolarizing TMV showing positively charged (magenta) and negatively charged (green) residues in VSD2. All four salt-bridge connections, ARG103-ASP49, ARG106-ASP49, ARG109-GLU59, and ARG109-ASP81, broke during the pulse and one new connection was formed: ARG106-ASP81. (b) An example showing how a Cl^-^ ion facilitates breakage of a salt bridge between ARG109 and GLU59 in VSD4 of NavMs. Other residues are omitted from representation for clarity. Frames were taken from the second simulation under hyperpolarizing TMV.

## Discussion

### Formation of complex pores is one explanation for the decrease in ionic current through VGICs observed experimentally

Electrophysiological measurements showed that electroporative submicrosecond electric pulses can cause a decrease in ionic currents through Nav, and Cav channels during action potentials in different excitable cells, whereby this decrease was suggested to involve electroconformational changes of the channels (6–8). Our simulations showed that electric fields can induce the formation of VSD pores that can expand into lipid-stabilized complex pores, leading to unfolding of the VSD from the channel. We expect that such unfolded VSDs become dysfunctional and unable to regulate the opening/closure of the channel pore. A dysfunctional ion channel cannot respond to changes in TMV anymore, which in electrophysiological measurements can indeed be detected as reduced current during action potentials. The complex pores (and the channel pore, if the channel got stuck in an intermediate open state) can only contribute to the “leak” current, which would be interpreted as the current flowing through lipid pores or other permeable defects caused by pulsed electric field. Overall, our simulations offer a mechanism that can potentially explain the experimentally measured decrease in ion channels’ currents.

Our simulations also revealed that, depending on the hydration and electrostatic properties, some VSDs can be more easily porated than others. This offers a possible explanation of why some channels appear to be affected at weaker electric fields than others (8). Furthermore, our simulations showed that some VSDs are more easily porated at hyperpolarizing (or depolarizing) TMV, which offers a possible mechanism of how the decrease in channel conductance depends on the polarity of the TMV (9). As different species express channels with different sequences, we anticipate that the electric-field mediated effects on ion channels should to some extent be species and also cell-type specific.

### VSD-associated complex pores: A type of long-lived metastable pores?

The mechanisms by which cell membranes remain permeable for seconds or minutes after application of electric pulses are still being discussed (4). Lipid pores do not seem to be stable long enough to account for this long-lived permeability as suggested based on their average lifetime: several μs as measured experimentally on pure lipid bilayers (59, 60), or few tens to hundreds of ns in the absence of external electric fields as estimated using MD (42, 61). It has been proposed that lipid pores could evolve into metastable pores involving both lipids and membrane proteins (47, 48). The complex pores observed in our simulations are indeed stabilized by lipid head-groups and VSD helices and have a long lifetime: after turning off the electric field we did not observe the pore closure within a 1-μs long simulation. This is longer than reported for closure of lipid pores in MD simulations (42, 61). Although we cannot infer from our simulations whether such complex pores could remain open for seconds or minutes after the pulse, it seems reasonable to anticipate that if a VSD unfolds to a major extent, it would not spontaneously refold back, thus leaving a long-lived permeable defect in the membrane until protein internalization by cell’s quality control machinery (62). Recent electrophysiological measurements indeed showed that, after exposing HEK cells to 300 ns pulses, there was about two-fold greater increase in the prolonged membrane conductance after the pulse in cells which expressed Cav channels compared to cells which did not (63). This could be due to Cav channels acting as sites of poration. It is further noteworthy that formation and ionic conduction of complex pores can be asymmetric with respect to TMV polarity. Such or similar protein-associated complex pores could therefore contribute to the not well understood mechanism of asymmetric cell membrane electropermeabilization, whereby exposure to electric pulses makes the membrane more extensively permeabilized either on anodic (hyperpolarized) or cathodic (depolarized) side (64–66).

Although our simulations mimicked exposure to high-intensity submicrosecond electric pulses, we expect that similar perturbation of VGICs could also occur during exposure to longer, more conventional microsecond or millisecond pulses with lower intensity. In one simulation of the mutant HCN1 channel under two-fold lower TMV of ∼0.7 V (this simulation was initially intended for a different study (52)), we observed formation of a complex pore at ∼5 μs after the onset of the electric field (Fig. S20). Thus we expect that VSD pores and complex pores can form also at lower TMV, just at a lower rate, similarly as observed for lipid pores (42).

### Activation of Nav channels by nanosecond pulsed electric fields

Depolarization of the membrane of excitable cells by external electric fields is known to trigger action potentials and is used for various purposes, for example artificial pacemaking, heart defibrillation, deep brain stimulation, and functional electrical stimulation. Stimulation of peripheral nerves is also an expected, though undesired, side effect in electroporation-based treatments which utilize microsecond or millisecond long pulses (67). High-intensity pulses with nano- and picosecond duration can activate VGICs and trigger action potentials as well (15, 68, 69). However, it remains unclear whether such short pulses are sufficiently long to activate VGICs by moving the VSDs during the pulse, or the activation is caused by post-pulse membrane depolarization resulting from ionic leakage across permeabilized membrane that disrupts the transmembrane ionic gradients. Stimulation of excitable cells by nanosecond pulses has been characterized by optical monitoring as the increase of intracellular Ca^2+^, and has been observed in bovine chromaffin cells exposed to 5 ns, 50 kV/cm pulse (16), adult rat cardiomyocytes exposed to 4 ns, 20-80 kV/cm pulses (15), and rat embryonic cardiomyocytes exposed to 10 ns, ≥36 kV/cm pulse (17). The first study (16) suggested that Ca^2+^ entered the cells through Cav channels, which were activated indirectly by post-pulse membrane depolarization. The second study (15) also suggested that Nav and Cav channels became activated by post-pulse membrane depolarization. Finally, the third study (17) reported that Cav-mediated increase in intracellular Ca^2+^ could be observed below the threshold for detectable membrane electropermeabilization (63 kV/cm). More recently, action potential generation in response to 200 ns, ≥1.9 kV/cm has been monitored in E18 rat hippocampal neurons by voltage-sensitive fluorescent dye (70). This study reported that action potentials could only be triggered above electropermeabilization threshold; however, it was possible that Nav channels were activated directly by the electric field. Direct activation of Nav channels by 12 ns pulses, not mediated by electropermeabilization, was also suggested in experiments on a peripheral nerve (12).

Activation of a voltage gated ion channel involves the movement of S4 helix, during which the positively charged residues on S4 break existing connections and form new ones with negatively charged residues on S1-S3 helices. Our analysis revealed that a high TMV, which can build on the membrane during application of electroporative pulses, can induce breakage and formation of salt bridges in the VSD on a nanosecond time scale. However, this occurs quite randomly in different VSDs of the channel and mainly involves formation of just one new salt bridge. An additional in-depth study is required to confirm or exclude the possibility that the TMV induced by high-intensity nanosecond pulses can activate VGICs.

## Conclusions

We performed MD simulations of three different VGICs under conditions mimicking exposure to pulsed electric fields. The following conclusions can be drawn from our results. Pulsed electric fields create conductive pores in the voltage-sensitive domains (VSDs) of voltage-gated ion channels, which can evolve into complex pores stabilized by lipid headgroups accompanied by unfolding of the VSD from the channel. VSDs are more or less prone to poration, depending on their hydration and electrostatic profile: the more hydrated the VSD is, and the more electrostatically favorable for the entry of ions, the more easily it porates. Poration of VSDs is one explanation for the decreased conductance of Nav and Cav channels observed in experiments. The differences in poration of different VSDs could explain why some channels are affected at lower pulse amplitude than others. Moreover, formation of VSD pores and complex pores, as well as their ionic conduction, can depend on the polarity of the transmembrane voltage. Hence, such pores could participate in the not completely understood asymmetric electropermeabilization of the cell membranes with respect to its anodic and cathodic side. We speculate that VSD-associated complex pores that lead to unfolding of the VSD from the protein, could act as long-lived permeable defects allowing enhanced transport of species during the seconds- or minutes-long cell membrane resealing.

Our study opens several new research directions. First, it will be interesting to investigate whether our findings are translatable to other membrane proteins. Our conclusions are mainly built on the analyses of protein hydration and electrostatic profiles, which can be straightforwardly performed on other membrane proteins. In this regard, a recent MD study on a human aquaporin has revealed formation of a transient electric-field induced pore through this protein; unlike the complex pores observed in our simulations, this pore closed within about 20 ns after turning off the electric field (71, 72). In our work we additionally reported conduction of ions through the central channel pore in the NavMs structure, which was presumably solved in an open state. Another possible research direction therefore would be to investigate electroporation of the different functional states of this and other channels. Finally, an interesting observation from our study is that in some cases, particularly in HCN1, lipid pores formed in addition to, or instead of, VSD pores. It is known that different lipids are differently suspectable for electroporation; for example, formation of lipid pores is facilitated in lipids with shorter tails or in peroxidized lipids (73, 74). Therefore, it will be interesting to investigate how the lipid environment affects poration of VGICs and other membrane proteins in upcoming studies.

## Author Contributions

L.R., M.K., I.T. and L.D. designed the research; L.R. performed simulations and analyzed data; L.R., M.K., I.T. and L.D. interpreted the results; L.R. wrote the manuscript, M.K., L.D. and I. T. revised the manuscript.

## Acknowledgements

This work was supported by grants from the Science for Life Laboratory and a synergy postdoc grant to L.D. and I.T. from KTH. The simulations were performed on resources provided by the Swedish National Infrastructure for Computing (SNIC) at PDC Centre for High Performance Computing (PDC-HPC). The authors thank Koushik Choudhury, Sergio Perez Conesa, and Damijan Miklavčič for useful comments to the manuscript.

## References

1. Yarmush, M.L., A. Golberg, G. Serša, T. Kotnik, and D. Miklavčič. 2014. Electroporation-Based Technologies for Medicine: Principles, Applications, and Challenges. Annu. Rev. Biomed. Eng. 16: 295–320.

2. Tieleman, D.P. 2004. The molecular basis of electroporation. BMC Biochem. 5: 1–12.

3. Delemotte, L., and M. Tarek. 2012. Molecular Dynamics Simulations of Lipid Membrane Electroporation. J. Membr. Biol. 245: 531–543.

4. Kotnik, T., L. Rems, M. Tarek, and D. Miklavčič. 2019. Membrane Electroporation and Electropermeabilization: Mechanisms and Models. Annu. Rev. Biophys. 48: 63–91.

5. Bezanilla, F. 2008. How membrane proteins sense voltage. Nat. Rev. Mol. Cell Biol. 9: 323–332.

6. Nesin, V., A.M. Bowman, S. Xiao, and A.G. Pakhomov. 2012. Cell permeabilization and inhibition of voltage-gated Ca2+ and Na+ channel currents by nanosecond pulsed electric field. Bioelectromagnetics. 33: 394–404.

7. Nesin, V., and A.G. Pakhomov. 2012. Inhibition of voltage-gated Na+ current by nanosecond pulsed electric field (nsPEF) is not mediated by Na+ influx or Ca2+ signaling. Bioelectromagnetics. 33: 443–451.

8. Yang, L., G.L. Craviso, P.T. Vernier, I. Chatterjee, and N. Leblanc. 2017. Nanosecond electric pulses differentially affect inward and outward currents in patch clamped adrenal chromaffin cells. PLoS One. 12: 1–20.

9. Chen, W., Y. Han, Y. Chen, and D. Astumian. 1998. Electric field-induced functional reductions in the K+ channels mainly resulted from supramembrane potential-mediated electroconformational changes. Biophys. J. 75: 196–206.

10. Chen, W., Z. Zhongsheng, and R.C. Lee. 2006. Supramembrane potential-induced electroconformational changes in sodium channel proteins: A potential mechanism involved in electric injury. Burns. 32: 52–59.

11. Pakhomov, A.G., I. Semenov, M. Casciola, and S. Xiao. 2017. Neuronal excitation and permeabilization by 200-ns pulsed electric field: An optical membrane potential study with FluoVolt dye. Biochim. Biophys. Acta - Biomembr. 1859: 1273–1281.

12. Casciola, M., S. Xiao, and A.G. Pakhomov. 2017. Damage-free peripheral nerve stimulation by 12-ns pulsed electric field. Sci. Rep. 7: 1–8.

13. Burke, R.C., S.M. Bardet, L. Carr, S. Romanenko, D. Arnaud-Cormos, P. Leveque, and R.P. O’Connor. 2017. Nanosecond pulsed electric fields depolarize transmembrane potential via voltage-gated K+, Ca2+ and TRPM8 channels in U87 glioblastoma cells. Biochim. Biophys. Acta - Biomembr. 1859: 2040–2050.

14. Dermol-Černe, J., D. Miklavčič, M. Reberšek, P. Mekuč, S.M. Bardet, R. Burke, D. Arnaud-Cormos, P. Leveque, and R. O’Connor. 2018. Plasma membrane depolarization and permeabilization due to electric pulses in cell lines of different excitability. Bioelectrochemistry. 122: 103–114.

15. Wang, S., J. Chen, M.T. Chen, P.T. Vernier, M.A. Gundersen, and M. Valderrábano. 2009. Cardiac myocyte excitation by ultrashort high-field pulses. Biophys. J. 96: 1640–1648.

16. Craviso, G.L., S. Choe, P. Chatterjee, I. Chatterjee, and P.T. Vernier. 2010. Nanosecond electric pulses: A novel stimulus for triggering Ca2+ influx into chromaffin cells via voltage-gated Ca2+ channels. Cell. Mol. Neurobiol. 30: 1259–1265.

17. Semenov, I., C. Zemlin, O.N. Pakhomova, S. Xiao, and A.G. Pakhomov. 2015. Diffuse, non-polar electropermeabilization and reduced propidium uptake distinguish the effect of nanosecond electric pulses. Biochim. Biophys. Acta - Biomembr. 1848: 2118–2125.

18. Blackiston, D.J., K.A. McLaughlin, and M. Levin. 2009. Bioelectric controls of cell proliferation: Ion channels, membrane voltage and the cell cycle. Cell Cycle. 8: 3527–3536.

19. Thrivikraman, G., S.K. Boda, and B. Basu. 2018. Unraveling the mechanistic effects of electric field stimulation towards directing stem cell fate and function: A tissue engineering perspective. Biomaterials. 150: 60–86.

20. Sula, A., J. Booker, L.C.T. Ng, C.E. Naylor, P.G. Decaen, and B.A. Wallace. 2017. The complete structure of an activated open sodium channel. Nat. Commun. 8: 14205.

21. Shen, H., Q. Zhou, X. Pan, Z. Li, J. Wu, and N. Yan. 2017. Structure of a eukaryotic voltage-gated sodium channel at near-atomic resolution. Science (80-.). 355: eaal4326.

22. Scheuer, T. 2014. Bacterial Sodium Channels: Models for Eukaryotic Sodium and Calcium Channels. Springer, Berlin, Heidelberg. pp. 269–291.

23. Catterall, W.A., and N. Zheng. 2015. Deciphering voltage-gated Na+ and Ca2+ channels by studying prokaryotic ancestors. Trends Biochem. Sci. 40: 526–534.

24. Lee, C.H., and R. MacKinnon. 2017. Structures of the Human HCN1 Hyperpolarization-Activated Channel. Cell. 168: 111–120.

25. Maor, E., A. Sugrue, C. Witt, V.R. Vaidya, C. V. DeSimone, S.J. Asirvatham, and S. Kapa. 2019. Pulsed electric fields for cardiac ablation and beyond: A state-of-the-art review. Hear. Rhythm. 16: 1112–1120.

26. Jo, S., T. Kim, V.G. Iyer, and W. Im. 2008. CHARMM-GUI: a web-based graphical user interface for CHARMM. J. Comput. Chem. 29: 1859–65.

27. Klauda, J.B., R.M. Venable, J.A. Freites, J.W. O’Connor, D.J. Tobias, C. Mondragon-Ramirez, I. Vorobyov, A.D. MacKerell, and R.W. Pastor. 2010. Update of the CHARMM All-Atom Additive Force Field for Lipids: Validation on Six Lipid Types. J. Phys. Chem. B. 114: 7830–7843.

28. Best, R.B., X. Zhu, J. Shim, P.E.M. Lopes, J. Mittal, M. Feig, and A.D. MacKerell. 2012. Optimization of the additive CHARMM all-atom protein force field targeting improved sampling of the backbone φ, ψ and side-chain χ1 and χ2 Dihedral Angles. J. Chem. Theory Comput. 8: 3257–3273.

29. Jorgensen, W.L., J. Chandrasekhar, J.D. Madura, R.W. Impey, and M.L. Klein. 1983. Comparison of simple potential functions for simulating liquid water. J. Chem. Phys. 79: 926–935.

30. Abraham, M.J., T. Murtola, R. Schulz, S. Páll, J.C. Smith, B. Hess, and E. Lindahl. 2015. GROMACS: High performance molecular simulations through multi-level parallelism from laptops to supercomputers. SoftwareX. 1–2: 19–25.

31. Páll, S., M.J. Abraham, C. Kutzner, B. Hess, and E. Lindahl. 2015. Tackling Exascale Software Challenges in Molecular Dynamics Simulations with GROMACS. Springer, Cham. pp. 3–27.

32. Nosé, S. 1984. A molecular dynamics method for simulations in the canonical ensemble. Mol. Phys. 52: 255–268.

33. Hoover, W.G. 1985. Canonical dynamics: Equilibrium phase-space distributions. Phys. Rev. A. 31: 1695–1697.

34. Parrinello, M., and A. Rahman. 1981. Polymorphic transitions in single crystals: A new molecular dynamics method. J. Appl. Phys. 52: 7182–7190.

35. Nosé, S., and M.L. Klein. 1983. Constant pressure molecular dynamics for molecular systems. Mol. Phys. 50: 1055–1076.

36. Darden, T., D. York, and L. Pedersen. 1993. Particle mesh Ewald: An *N* ·log(*N*) method for Ewald sums in large systems. J. Chem. Phys. 98: 10089–10092.

37. Hess, B., H. Bekker, H.J.C. Berendsen, and J.G.E.M. Fraaije. 1997. LINCS: A linear constraint solver for molecular simulations. J. Comput. Chem. 18: 1463–1472.

38. Casciola, M., M.A. Kasimova, L. Rems, S. Zullino, F. Apollonio, and M. Tarek. 2016. Properties of lipid electropores I: Molecular dynamics simulations of stabilized pores by constant charge imbalance. Bioelectrochemistry. 109: 108–116.

39. Kutzner, C., H. Grubmüller, B.L. De Groot, and U. Zachariae. 2011. Computational electrophysiology: The molecular dynamics of ion channel permeation and selectivity in atomistic detail. Biophys. J. 101: 809–817.

40. Humphrey, W., A. Dalke, and K. Schulten. 1996. VMD: Visual molecular dynamics. J. Mol. Graph. 14: 33–38.

41. Aksimentiev, A., and K. Schulten. 2005. Imaging α-Hemolysin with Molecular Dynamics: Ionic Conductance, Osmotic Permeability, and the Electrostatic Potential Map. Biophys. J. 88: 3745–3761.

42. Levine, Z.A., and P.T. Vernier. 2010. Life Cycle of an Electropore: Field-Dependent and Field-Independent Steps in Pore Creation and Annihilation. J. Membr. Biol. 236: 27–36.

43. Böckmann, R.A., B.L. de Groot, S. Kakorin, E. Neumann, and H. Grubmüller. 2008. Kinetics, statistics, and energetics of lipid membrane electroporation studied by molecular dynamics simulations. Biophys. J. 95: 1837–50.

44. Hibino, M., M. Shigemori, H. Itoh, K. Nagayama, and K. Kinosita. 1991. Membrane conductance of an electroporated cell analyzed by submicrosecond imaging of transmembrane potential. Biophys. J. 59: 209–220.

45. Frey, W., J.A. White, R.O. Price, P.F. Blackmore, R.P. Joshi, R. Nuccitelli, S.J. Beebe, K.H. Schoenbach, and J.F. Kolb. 2006. Plasma membrane voltage changes during nanosecond pulsed electric field exposure. Biophys. J. 90: 3608–15.

46. Benz, R., and F. Conti. 1981. Reversible electrical breakdown of squid giant axon membrane. Biochim. Biophys. Acta - Biomembr. 645: 115–123.

47. Weaver, J.C., and P.T. Vernier. 2017. Pore lifetimes in cell electroporation: Complex dark pores? arXiv. 1708.07478, http://arxiv.org/abs/1708.07478.

48. Weaver, J.C., and A. Barnett. 2012. Progress toward a Theoretical Model for Electroporation Mechanism: Membrane Electrical Behavior and Molecular Transport. In: Chang DC, BM Chassy, JA Saunders, AE Sowers, editors. Guide to Electroporation and Electrofusion. Academic Press. pp. 91–117.

49. Ulmschneider, M.B., C. Bagnéris, E.C. McCusker, P.G. Decaen, M. Delling, D.E. Clapham, J.P. Ulmschneider, and B.A. Wallace. 2013. Molecular dynamics of ion transport through the open conformation of a bacterial voltage-gated sodium channel. Proc. Natl. Acad. Sci. U. S. A. 110: 6364–9.

50. Tokman, M., J.H. Lee, Z.A. Levine, M.-C. Ho, M.E. Colvin, and P.T. Vernier. 2013. Electric Field-Driven Water Dipoles: Nanoscale Architecture of Electroporation. PLoS One. 8: e61111.

51. Sun, S., J.T.Y. Wong, and T.-Y. Zhang. 2011. Atomistic Simulations of Electroporation in Water Preembedded Membranes. J. Phys. Chem. B. 115: 13355–13359.

52. Kasimova, M.A., D. Tewari, J. Cowgill, W. Carrasquel Ursulaez, J. Lin, L. Delemotte, and B. Chanda. 2019. Helix Breaking Transition in the S4 of HCN Channel is Critical for Hyperpolarization-Dependent Gating. SSRN Electron. J.: http://dx.doi.org/10.2139/ssrn.3468492.

53. Ho, M.-C., M. Casciola, Z.A. Levine, and P.T. Vernier. 2013. Molecular Dynamics Simulations of Ion Conductance in Field-Stabilized Nanoscale Lipid Electropores. J. Phys. Chem. B. 117: 11633–11640.

54. Casciola, M., M.A. Kasimova, L. Rems, S. Zullino, F. Apollonio, and M. Tarek. 2016. Properties of lipid electropores I: Molecular dynamics simulations of stabilized pores by constant charge imbalance. Bioelectrochemistry. 109: 108–116.

55. Rems, L., M. Tarek, M. Casciola, and D. Miklavčič. 2016. Properties of lipid electropores II: Comparison of continuum-level modeling of pore conductance to molecular dynamics simulations. Bioelectrochemistry. 112: 112–124.

56. Delemotte, L., M. Tarek, M.L. Klein, C. Amaral, and W. Treptow. 2011. Intermediate states of the Kv1.2 voltage sensor from atomistic molecular dynamics simulations. Proc. Natl. Acad. Sci. U. S. A. 108: 6109–6114.

57. Amaral, C., V. Carnevale, M.L. Klein, and W. Treptow. 2012. Exploring conformational states of the bacterial voltage-gated sodium channel NavAb via molecular dynamics simulations. Proc. Natl. Acad. Sci. U. S. A. 109: 21336–21341.

58. Kasimova, M.A., E. Lindahl, and L. Delemotte. 2018. Determining the molecular basis of voltage sensitivity in membrane proteins. J. Gen. Physiol. 215: 1444–1458.

59. Melikov, K.C., V.A. Frolov, A. Shcherbakov, A. V. Samsonov, Y.A. Chizmadzhev, and L. V. Chernomordik. 2001. Voltage-Induced Nonconductive Pre-Pores and Metastable Single Pores in Unmodified Planar Lipid Bilayer. Biophys. J. 80: 1829–1836.

60. Sengel, J.T., and M.I. Wallace. 2016. Imaging the dynamics of individual electropores. Proc. Natl. Acad. Sci. U. S. A. 113: 5281–6.

61. Bennett, W.F.D., N. Sapay, and D.P. Tieleman. 2014. Atomistic Simulations of Pore Formation and Closure in Lipid Bilayers. Biophys. J. 106: 210–219.

62. Grant, B.D., and J.G. Donaldson. 2009. Pathways and mechanisms of endocytic recycling. Nat. Rev. Mol. Cell Biol. 10: 597–608.

63. Hristov, K., U. Mangalanathan, M. Casciola, O.N. Pakhomova, and A.G. Pakhomov. 2018. Expression of voltage-gated calcium channels augments cell susceptibility to membrane disruption by nanosecond pulsed electric field. Biochim. Biophys. Acta - Biomembr. 1860: 2175–2183.

64. Hibino, M., H. Itoh, K. Kinosita, and Jr. 1993. Time courses of cell electroporation as revealed by submicrosecond imaging of transmembrane potential. Biophys. J. 64: 1789–800.

65. Krassen, H., U. Pliquett, and E. Neumann. 2007. Nonlinear current–voltage relationship of the plasma membrane of single CHO cells. Bioelectrochemistry. 70: 71–77.

66. Gabriel, B., and J. Teissié. 1997. Direct observation in the millisecond time range of fluorescent molecule asymmetrical interaction with the electropermeabilized cell membrane. Biophys. J. 73: 2630–2637.

67. Miklavčič, D., B. Mali, B. Kos, R. Heller, and G. Serša. 2014. Electrochemotherapy: from the drawing board into medical practice. Biomed. Eng. Online. 13: 29.

68. Vernier, P.T., Y. Sun, M.-T. Chen, M.A. Gundersen, and G.L. Craviso. 2008. Nanosecond electric pulse-induced calcium entry into chromaffin cells. Bioelectrochemistry. 73: 1–4.

69. Semenov, I., S. Xiao, D. Kang, K.H. Schoenbach, and A.G. Pakhomov. 2015. Cell stimulation and calcium mobilization by picosecond electric pulses. Bioelectrochemistry. 105: 65–71.

70. Pakhomov, A.G., I. Semenov, M. Casciola, and S. Xiao. 2017. Neuronal excitation and permeabilization by 200-ns pulsed electric field: An optical membrane potential study with FluoVolt dye. Biochim. Biophys. Acta - Biomembr. 1859: 1273–1281.

71. Marracino, P., M. Bernardi, M. Liberti, F. Del Signore, E. Trapani, J.A. Gárate, C.J. Burnham, F. Apollonio, and N.J. English. 2018. Transprotein-Electropore Characterization: A Molecular Dynamics Investigation on Human AQP4. ACS Omega. 3: 15361–15369.

72. Bernardi, M., P. Marracino, M. Liberti, J.A. Gárate, C.J. Burnham, F. Apollonio, and N.J. English. 2019. Controlling ionic conductivity through transprotein electropores in human aquaporin 4: A non-equilibrium molecular-dynamics study. Phys. Chem. Chem. Phys. 21: 3339–3346.

73. Gurtovenko, A.A., and A.S. Lyulina. 2014. Electroporation of Asymmetric Phospholipid Membranes. J. Phys. Chem. B. 118: 9909–9918.

74. Vernier, P.T., Z.A. Levine, Y.-H. Wu, V. Joubert, M.J. Ziegler, L.M. Mir, and D.P. Tieleman. 2009. Electroporating Fields Target Oxidatively Damaged Areas in the Cell Membrane. PLoS One. 4: e7966.

